# Mutations in *emb*B406 are associated with low-level ethambutol resistance in Canadian *Mycobacterium tuberculosis* isolates

**DOI:** 10.1101/2023.03.08.531832

**Authors:** Morgan R. Hiebert, Meenu K. Sharma, Melissa J. Rabb, Lisa J. Karlowsky, Kiana S. Bergman, Hafid Soualhine

**Affiliations:** National Reference Centre for Mycobacteriology, National Microbiology Laboratory, Public Health Agency of Canada, Winnipeg, MB, Canada; Department of Medical Microbiology and Infectious Diseases, University of Manitoba, Winnipeg, MB, Canada

**Keywords:** ethambutol, resistance, *emb*B, *Mycobacterium tuberculosis*

## Abstract

**Background:** In *Mycobacterium tuberculosis*, molecular predictions of ethambutol resistance rely primarily on the detection of mutations within *emb*B. However, discordance between *emb*B406 mutations and phenotypic drug sensitivity questions its clinical significance. This study aims to decipher the association of *emb*B406 mutations with ethambutol resistance in *M. tuberculosis*.

**Methods:** All *M. tuberculosis* isolates from our culture collection containing *emb*B406 mutations (n=16) and pan-sensitive control isolates (n=10) were selected for this study. Phenotypic drug susceptibility testing for ethambutol was performed in duplicate on the BACTEC™ MGIT™ 960 at concentrations of 2, 3, 4, and 5 μg/mL with strain H37Rv as assay control. Whole genome sequencing was performed on Illumina Miseq for drug resistance predictions (MyKrobe Predictor v.0.7.0), phylogenomics (SNVPhyl v.1.2.3) and single nucleotide polymorphism analysis (Snippy).

**Results:** Two *emb*B406 mutation subtypes were found among 16 strains: Gly406Asp and Gly406Ala. MyKrobe predicted all strains of either subtype to be ethambutol resistant. However, 12 of 16 strains appear phenotypically sensitive at 5 μg/mL but exhibit variable resistance between 2-4 μg/mL. Of these 12 strains, a newly described frameshift mutation in regulator *embR* (Gln258fs) was found in 9 strains.

**Conclusions:** Mutations in *emb*B406 are associated with low-level ethambutol resistance currently undetectable by the critical concentration of 5 μg/mL for ethambutol. Novel mutations are predicted to exacerbate variability in ethambutol resistance. We suggest amendment to molecular and phenotypic drug susceptibility testing to improve ethambutol DST sensitivity and specificity as well as concordance between rapid and gold standard methods.

## Introduction

*Mycobacterium tuberculosis* remains a priority pathogen for the World Health Organization (WHO) as the causative agent of tuberculosis (TB) and the second leading infectious cause of mortality worldwide (1). The rise of antimicrobial resistance is a concern, as treatment options are limited for resistant cases (2–4). Globally, multi-drug resistant (MDR) TB poses a serious risk to public health (1). In Canada, nearly 10% of all TB cases from 2017 to 2020 were resistant to one or more first-line anti-tuberculosis drugs (5).

Drug susceptibility testing (DST) for *M. tuberculosis* infections includes culture-based methods as the gold standard (6). However, these phenotypic assays are time-consuming due to the slow growing nature of *M. tuberculosis*. Advances in molecular techniques such as PCR-based methods (7–10) and whole genome sequencing (11–13) (WGS) have enabled rapid detection of resistance-associated gene mutations. However, discordance between gold standard phenotypic DST and rapid molecular methods has been observed in some cases (14–16).

Ethambutol is a bacteriostatic first-line anti-tuberculosis drug that targets arabinosyltransferases encoded by the *emb*CAB operon, hindering the arabinogalactan biosynthetic pathway and thereby inhibiting mycobacterial cell wall synthesis (17). Mutations in the arabinosyltransferase-encoding *emb*B gene, including codon 406, are known to convey ethambutol resistance in *M. tuberculosis* and are employed in diagnostic pipelines for rapidly predicting resistance to ethambutol (18, 19). However, several studies have reported *emb*B406 mutations in both ethambutol resistant and susceptible samples (15, 16, 20–22). This discordance between genotypic and phenotypic DST at the current critical concentration questions the clinical impact of *emb*B406 mutations on the efficacy of treatment regimens that include ethambutol.

We predict that *emb*B mutations in codon 406 are associated with low-level ethambutol resistance which is undetectable by the current critical concentration (5 μg/mL) utilized for DST on the BACTEC™ MGIT™ 960 system. This study evaluates the impact of specific *emb*B406 mutations on ethambutol resistance in Canadian *M. tuberculosis* isolates and tabulates novel mutations outside of *emb*B that may exacerbate phenotypic variability.

## Methods

### Study Strains

The culture collection housed within the National Reference Centre for Mycobacteriology (2003-2022) was screened for *M. tuberculosis* isolates containing *emb*B406 mutations. Screening by routine sequencing of the *emb*B gene (described below) identified 16 isolates harboring an *emb*B406 mutation. A total of 10 pan-sensitive isolates lacking *emb*B mutations were selected as controls in addition to *M. tuberculosis* strain H37Rv ATCC 27294.

### DNA Extraction

Genomic DNA was extracted from isolates using the DNeasy® UltraClean® Microbial Kit (Qiagen, Germany) with the following modifications to the manufacturer’s instructions. A loopful of solid culture grown on Middlebrook 7H10 media (Becton, Dickinson and Company, NJ) was added to PowerBead solution in a PowerBead tube and vortexed for three seconds before boiling for 10 minutes to ensure non-viability. Following the addition of Solution SL, suspensions were bead-beaten in a FastPrep-24 tissue homogenizer (MP Biomedicals, CA) for 3 cycles (4 m/s for 30 s; 1 min rest on ice between cycles). Extracted genomic DNA was quantified on a Qubit® 3.0 fluorometer (Invitrogen, MA) with a dsDNA high sensitivity assay kit (Invitrogen, MA) and quality was assessed on a NanoDrop™ 2000 spectrophotometer (Thermofisher Scientific, MA).

### Sanger Sequencing and Data Analysis

The *emb*B region containing codon 406 was amplified using PCR primers F1 5’-TGATATTCGGCTTCCTGCTC-3’ and R1 5’-TGCACACCCAGTGTGAATG-3’ designed using Primer3 (23) (version 0.4.0) and acquired from ThermoFisher Scientific. Primers were utilized at 0.2 μM in Amplitaq Gold™ 360 Master Mix (Applied Biosystems, MA). PCR amplification programs consisted of a 10-minute initial denaturation at 95°C followed by 30 cycles (94°C, 30 sec; 60°C, 30sec; 72°, 30 sec) and a final extension at 72°C for 7 minutes. PCR products were purified using PCRClean™ DX (Aline Biosciences, MA) according to the manufacturer’s instructions (version Rev.2.10). Sanger sequencing was performed on a 3730xl DNA Analyzer (Applied Biosystems, MA) utilizing the same primers as for amplification. Sequence data from primers were paired and trimmed in SeqMan Pro 15 (DNASTAR, WI) before MUSCLE alignment (24) to the *M. tuberculosis* H37Rv *emb*B reference sequence (NC_000962.3: 4246514-4249810) in Geneious (version 11.0.12; www.geneious.com).

### Whole Genome Sequencing

DNA libraries were prepared with the Illumina® DNA Prep kit (Illumina, CA) using a modified quarter-volume protocol before sequencing on the MiSeq platform with a MiSeq Reagent Kit v3 (500-cycle; Illumina, CA) to generate 2×250 bp paired-end reads. Completed sequencing runs were uploaded to the Integrated Rapid Infectious Disease Analysis (IRIDA) Platform (25). Trimmomatic (26) in Galaxy (version 0.36.5) was utilized to trim sequences resulting from adaptor read-through (ILLUMINACLIP:Nextera-PE:2:30:10:8) and poor-quality reads (SLIDINGWINDOW:5:20). We assessed read quality with FastQC (27) (version 0.72). Kraken2 (version 2.2; github.com/DerrickWood/kraken2) was employed to detect potential contaminating microbial DNA. BioHansel (28) (version 1.2), an in-house developed tool, was utilized for *M. tuberculosis* complex differentiation. Using Snippy (github.com/tseemann/snippy), sequencing reads were assembled and compared to an annotated *M. tuberculosis* H37Rv reference genome (AL_123456.3) for single nucleotide polymorphism (SNP) calling. The quality of the alignment used for SNP detection was determined with QualiMap2 (29). Minimum quality thresholds for alignments included genome coverage of 30, 90% of reads mapping to the reference genome, a mapping quality of 58.5, and GC content of 65 ± 1%. Alignments were visualized in Geneious (version 11.0.12; www.geneious.com). MyKrobe Predictor (19) (version 0.7.0) was utilized for genotypic DST for first- and second-line anti-tuberculosis drugs as part of routine diagnostic workflows. SNVPhyl (30) (version 1.2.3) was used for the phylogenomic analysis of all strains in the study, utilizing the *M. tuberculosis* H37Rv genome as a reference (NC_000962.3).

### Phenotypic Drug Susceptibility Testing

DST for first line anti-tuberculosis drugs was performed on the BD BACTEC™ MGIT™ 960 automated mycobacterial detection system with the BD BBL™ MGIT™ AST SIRE kit (Becton, Dickinson and Company, NJ). All isolates were tested at the following critical concentrations as described by the manufacturer and the Clinical and Laboratory Standards Institute (CLSI) recommendations: 1.00 μg/mL rifampin, 0.1 μg/mL isoniazid, 100 μg/mL pyrazinamide, 5 μg/mL ethambutol (6). For ethambutol (Sigma-Aldrich, MO), additional concentrations of 4 μg/mL, 3 μg/mL, and 2 μg/mL were tested on the MGIT™ 960 system. The inoculum for DST was prepared from solid culture according to the manufacturer’s instructions (version E 2004/2006). Antibiotic susceptibility was determined using the proportion method, in which growth in drug-containing media was compared to that of the drug-free growth control, according to the manufacturer’s instructions (version E 2004/6).

Additionally, the inoculum purity was tested by subculturing to blood agar plates (Oxoid, ON) at 37°C for 48 hours. DST for all strains was performed in parallel with *M. tuberculosis* H37Rv ATCC27294 control strain. To ensure reproducibility, two different technicians each performed DST on every sample. Any discrepancy between two results was resolved by repeating DST a third time and accepting the majority result expressed by two of the three tests.

## Results

### Genotypic Susceptibility Predictions

The genotypic ethambutol susceptibility profiles of 26 *M. tuberculosis* strains were analyzed using *emb*B-targeted PCR-based sequencing and WGS (Figure 1). Of the 16 strains with *emb*B406 mutations, 13 strains harboured a Gly406Asp mutation and 3 strains exhibited a Gly406Ala mutation. The remaining 10 control strains retained a wild-type *emb*B406 locus. MyKrobe Predictor output indicated that all 16 strains with *emb*B406 mutations were predicted to be ethambutol resistant. As anticipated, all strains with a wild-type *emb*B406 locus were predicted to be sensitive to ethambutol.

**Figure 1.**
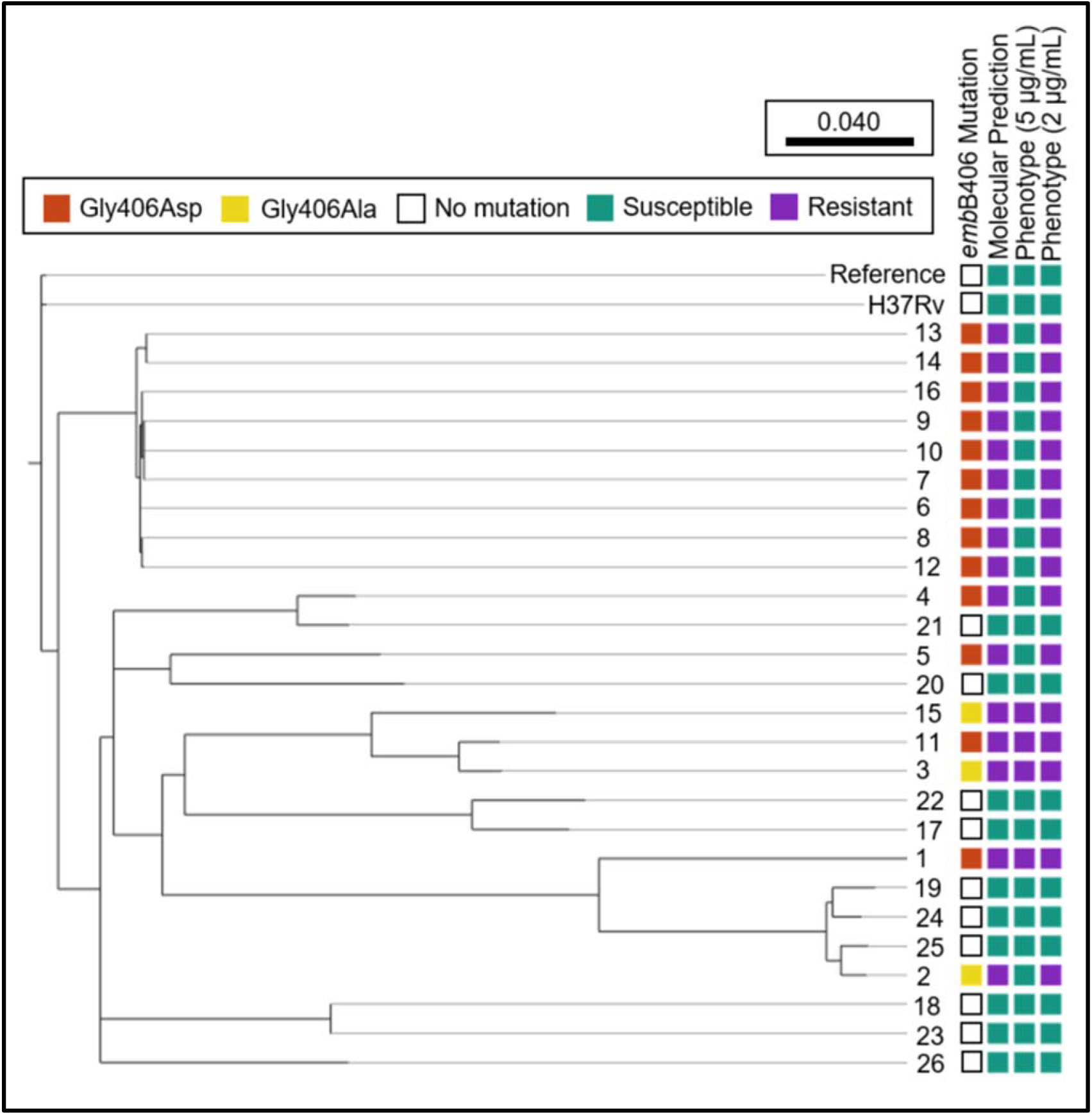
Ethambutol susceptibility predictions among *M. tuberculosis* strains harbouring *emb*B406 mutations. Depiction of phylogeny and antibiotic susceptibility metadata for study strains. Phylogenetic tree generated with SNVPhyl, visualized in Microreact. Antibiotic susceptibility prediction determined by MyKrobe Predictor. Ethambutol phenotype (resistant or susceptible) depicted at the current critical concentration, 5 μg/mL, and a lower hypothetical concentration, 2 μg/mL. Tree reference constitutes the literature *M. tuberculosis* H37Rv whole genome sequence (NC_000962.3) and the adjacent H37Rv which was cultured and sequenced in-house.

### Phenotypic Ethambutol Susceptibility

Ethambutol susceptibility of 26 *M. tuberculosis* strains was assessed by phenotypic DST on the BACTEC™ MGIT™ 960 system (Figure 1). All 10 control strains with a wild-type *emb*B406 locus were sensitive to ethambutol at all concentrations (2 μg/mL – 5 μg/mL). Notably, 4/16 (25%) strains with an *emb*B406 mutation were resistant to ethambutol at the critical concentration of 5 μg/mL. Of these 4 strains, 2 had a Gly406Asp mutation while the other 2 harboured a Gly406Ala mutation. By mutation subtype, 2/13 (15.4%) Gly406Asp strains and 2/3 (66.6%) Gly406Ala strains were resistant at 5 μg/mL.

The remaining 12/16 (75%) strains were susceptible to ethambutol at 5 μg/mL, however they exhibited resistance below the critical concentration (Table 1). Specifically, 2 strains were resistant at 4 μg/mL, 5 were resistant at 3 μg/mL, and 5 were resistant at 2 μg/mL.

**Table 1.**
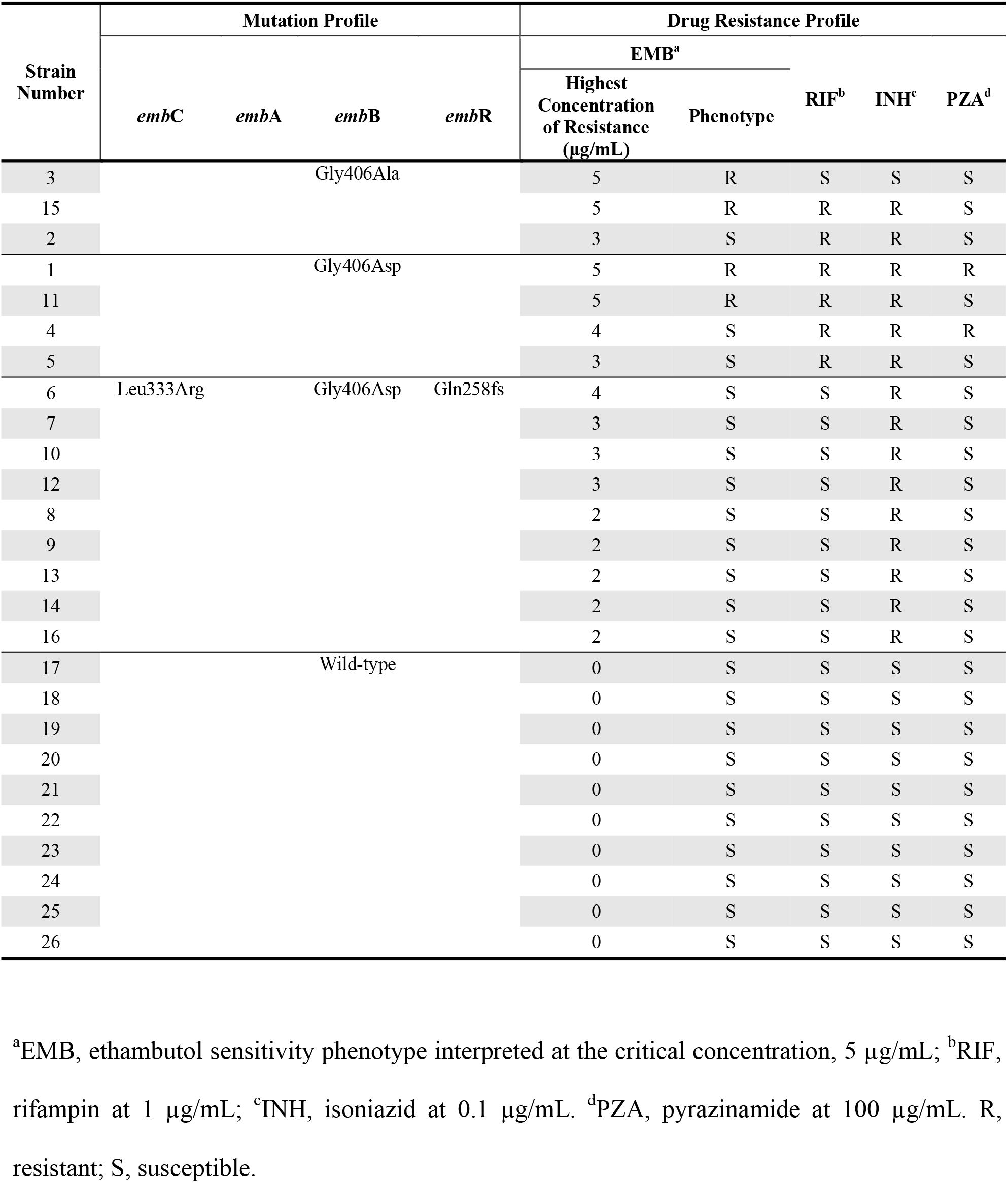
Drug resistance and *embCAB* mutation profiles of *M. tuberculosis* strains.

### *emb*CAB Mutation Profiles

Given the phenotypic variability of strains with identical *emb*B406 mutations, we surveyed additional mutations that may contribute to ethambutol resistance (≥ 2 μg/mL) by WGS. In particular, non-synonymous mutations within components of the *emb*CAB system were compiled, including operon regulator *embR* (Figure 2). Several novel mutations were found exclusively in ethambutol-resistant strains. In *embC*, a Leu333Arg substitution was found in 9 strains. One strain exhibited a Tyr502Cys substitution in *emb*A. In addition to the *emb*B406 mutations described above (Gly406Asp, n=13; Gly406Ala, n=3), one strain exhibited a Thr1082Ala substitution in the C-terminal domain of *emb*B. In *emb*R, 9 strains have a frameshift mutation in codon 258 (Gln258fs), located upstream from the forkhead-associated domain.

**Figure 2.**
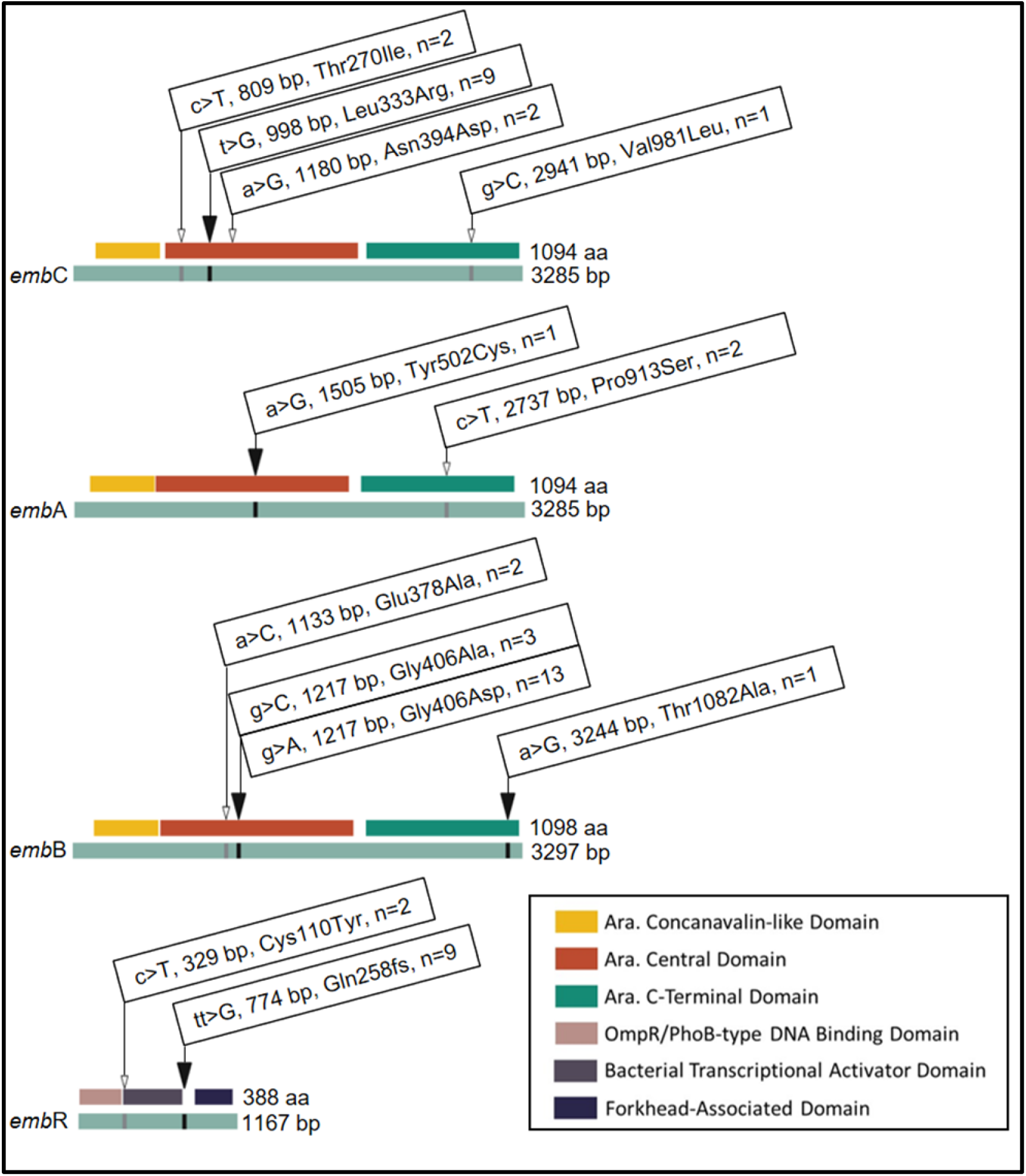
Non-synonymous mutations found in components of *embCAB* system of *M. tuberculosis* strains exhibiting ethambutol resistance. Linear gene (light blue) and protein domain (key) map of SNPs in the *emb*CAB operon. Identity and position of base pair (bp) and amino acid (aa) substitutions are provided. n, total number of EMB-resistant strains harbouring a given mutation. Black solid triangle, found in *emb*B406 mutants only. White triangle, found in both *emb*B406 mutants and pan-susceptible control strains.

All mutations observed exclusively in two or more ethambutol-resistant strains were organized into mutation profiles according to strain, DST phenotype, and maximum concentration of resistance (Table 1). Ultimately, this resulted in the identification of three mutation profiles: (A) *emb*B Gly406Asp-only, (B) *emb*B Gly406Ala-only, and (C) *emb*C Leu333Arg, *emb*B Gly406Asp, and *emb*R Gln258.

In general, strains that exclusively contained an *emb*B406 mutation exhibited resistance at maximum concentrations ranging from 3 μg/mL to 5 μg/mL. Notably, all 9 strains which held three identical mutations in *emb*C (Leu333Arg), *emb*B (Gly406Asp), and *emb*R (Gln258fs) exhibit ethambutol sensitivity according to the gold standard phenotypic DST at the critical concentration of 5 μg/ml. Strains exhibiting this mutation profile were resistant to ethambutol at maximum concentrations ranging from 2 μg/mL to 4 μg/mL.

### Phenotypic Susceptibility to Additional First Line Drugs

Phenotypic susceptibility to the remaining three first line drugs rifampin, isoniazid, and pyrazinamide was also assessed by critical concentration on the BACTEC™ MGIT™ 960 system (Table 1). All 10 control strains with a wild-type *emb*B406 locus were pan-sensitive to all first line drugs. According to mutation profile, all strains with *embC* Leu333Arg, *emb*B Gly406Asp, and *embR* Gln258 mutations were resistant to isoniazid while remaining sensitive to rifampin, pyrazinamide and ethambutol at the critical concentration. In contrast, 6/7 (85.7%) strains harbouring only an *emb*B406 mutation (Gly406Ala or Gly406Asp) were MDR, exhibiting resistance to both rifampin and isoniazid at the critical concentrations. Of these strains, 2/6 (33.3%) were resistant to all four first line drugs at the critical concentration (Gly406Asp, n=2), 2/6 (33.3%) were resistant to rifampin, isoniazid, and ethambutol (Gly406Asp, n=1; Gly406Ala, n=1), and 2/6 (33.3%) were resistant to rifampin and isoniazid only (Gly406Asp, n=1; Gly406Ala, n=1). A single isolate exhibited ethambutol mono-resistance (Gly406Ala).

## Discussion

According to the Canadian tuberculosis standards (31), and CLSI guidelines (6), routine phenotypic DST for ethambutol employs a critical concentration of 5 μg/mL on the BACTEC™ MGIT™ 960 system. While this method remains the gold standard, rapid sequencing-based approaches enable drug susceptibility predictions weeks before phenotypic susceptibility results are available. This requires comprehensive knowledge of molecular predictors of resistance to accurately inform treatment regimens. However, our recent experience with WGS-based molecular DST has detected discrepant phenotypic DST results in several isolates. This corroborates numerous studies highlighting discordance between ethambutol genotypic susceptibility predictions and phenotypic DST at a critical concentration of 5 μg/mL (14–16, 20–22).

The present study evaluates ethambutol resistance among *M. tuberculosis* strains with *emb*B406 mutations. While commonly employed in diagnostic pipelines for rapid DST (i.e., MyKrobe Predictor), *emb*B406 mutations are disputed as a marker for ethambutol resistance. Indeed, we report that 75% (12/16) of strains with *emb*B406 mutations in our culture collection were susceptible at the current critical concentration using the BACTEC™ MGIT™ 960 system. However, DST performed at 2, 3, and 4 μg/mL showed that all susceptible strains with *emb*B406 mutations exhibited ethambutol resistance undetectable by the current critical concentration. Stratifying by mutation subtype, 11/13 (84.6%) Gly206Asp strains and 1/3 (33.3%) Gly406Ala strains exhibited resistance below 5 μg/mL. Specifically, 2/13 (15.4%) Gly406Asp strains were resistant at 4 μg/mL, 4/13 (30.8%) Gly406Asp strains and 1/3 (33.3%) Gly406Ala strains were resistant at 3 μg/mL, and 5/13 (38.5%) Gly406Asp strains were resistant at 2 μg/mL. In our limited collection, Gly406Asp appears to be associated with low-level resistance.

Expanding our scope, we investigated other genes within the *emb*CAB operon, including the operon’s regulator *emb*R. As a result, we found that 9 strains exhibited identical mutations in *emb*C (Leu333Arg) and *emb*R (Gln258fs) which have not been previously described, in addition to the *emb*B Gly406Asp mutation. Interestingly, all 9 strains were sensitive at 5 μg/mL but exhibited low-level resistance undetectable by the critical concentration. Of these strains, 1/9 (11.1%) was resistant at 4 μg/mL, 3/9 (33.3%) were resistant at 3 μg/mL, and 5/9 (55.6%) were resistant at 2 μg/mL. Notably, the *emb*R frameshift mutation occurs within the forkhead-associated (FHA) domain of the operon’s regulator. As the FHA domain is critical for phosphorylation and activation of *emb*R (32), we predict that this novel and disruptive frameshift mutation may impact *emb*CAB expression and cause variability in resistance. Further investigation of these mutations is required to isolate their impact on ethambutol resistance.

Briefly, we investigated susceptibility to the remaining three first line anti-tuberculosis drugs: isoniazid, rifampin, and pyrazinamide. Ethambutol mono-resistance, while rare, was observed in a single strain with an *emb*B Gly406Ala mutation. Isoniazid resistance was observed in 15/16 (93.75%) of strains harbouring a mutation in *emb*B406, suggesting an association between ethambutol and isoniazid resistance. Previously, others have suggested common mechanisms of resistance exist between isoniazid and ethambutol given their shared drug target (33) and synergistic activity (34). Additional studies are required to solidify a mechanistic linkage between isoniazid and ethambutol resistance.

Explored at length above, discordance between molecular methods and phenotypic DST on the BACTEC™ MGIT™ 960 system raises questions to the current critical concentration for phenotypic DST as well as the clinical impact of *emb*B406 mutations on ethambutol resistance. However, these discrepancies are not isolated to the BACTEC™ MGIT™ 960 system but span to other methods detailed by the CLSI guidelines for susceptibility testing (6). DST by agar proportion on solid media employs varying critical concentrations for ethambutol depending on media type: 2 μg/mL by Löwenstein-Jensen (LJ), 5 μg/mL by Middlebrook 7H10, and 7.5 μg/mL by Middlebrook 7H11 (6). Few studies describe concordance between *emb*B406 mutations and phenotypic DST using the agar proportion method (Table 2) (35, 36). Several have found *emb*B406 mutation subtypes among both resistant and susceptible strains by agar proportion on LJ media as detailed in Table 2 (22, 37–39). Park *et al*. describe moderate ethambutol resistance (0.88 – 4 μg/mL) among *M. tuberculosis* isolates with *emb*B406 mutations (22). Others employing agar proportion DST have suggested that ethambutol resistance is multigenic and that *emb*B406 mutations along are insufficient for the development of resistance (20, 40).

**Table 2.**
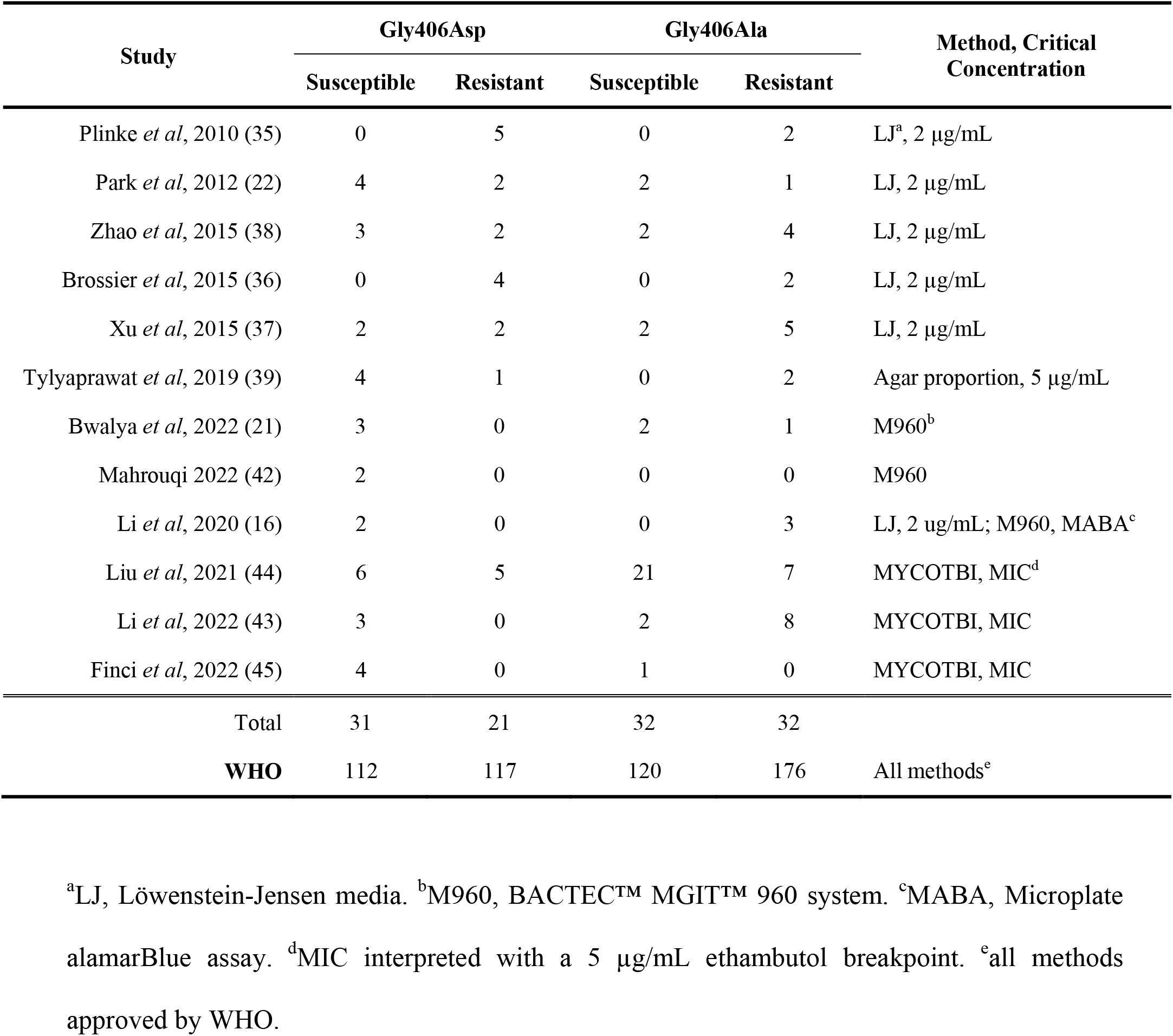
Ethambutol DST and *emb*B406 mutations among literature.

More recent commercial automated broth-based systems include the obsolete BACTEC™ MGIT™ 460 radiometric system and the current BACTEC™ MGIT™ 960 system employ agar proportion-equivalent ethambutol critical concentrations of 2.5 μg/mL and 5 μg/mL, respectively (6, 15). The adaptation of critical concentrations from agar proportion methods to the MGIT™ 460 system, which were shortly modified in the shift to the MGIT™ 960 system has been criticized for poor concordance and standardization (15, 41). Again, several describe ethambutol susceptible strains with *emb*B406 mutations (15, 16, 21, 42). Further, Bwalya *et al*. (2022) and Christianson *et al*. (2014) discuss low-level ethambutol resistance among strains with *emb*B406 mutations. These observations are accompanied by a recommendation that the BACTEC™ MGIT™ 960 critical concentration be lowered to ensure resistance at or near the 5 μg/mL critical concentration is detected (15).

Microdilution plates have sought to address inconsistencies between phenotypic DST methods by assessing minimum inhibitory concentrations (MICs) rather than critical concentrations for ethambutol(41). Studies employing the Sensititre™ *Mycobacterium tuberculosis* MYCOTBI AST plate also report *emb*B406 mutations among strains exhibiting resistance below the 5 μg/mL breakpoint for ethambutol(43–45). Liu *et al*. (2021) explain that *emb*B406 mutations elevated ethambutol MICs at sub-threshold levels (44). Li *et al*. (2022) record *emb*B406 variants with MICs ranging from 1 – 16 μg/mL (43).

Phenotypic DST methods and critical concentrations endorsed by the WHO include agar proportion, commercial broth systems, and broth microdilution plates (18). The 2021 WHO catalog of *Mycobacterium tuberculosis* complex mutations associated with drug resistance records 112 sensitive and 117 resistant strains with Gly406Asp mutations while detailing a positive predictive value (PPV) of 59.5%, specificity of 99.5%, and sensitivity of 3.6% for ethambutol phenotypic DST (18). In addition, the WHO reports 120 sensitive and 176 resistant strains with Gly406Ala mutations, recording a PPV of 51.1%, specificity of 99.6%, and sensitivity of 2.4% for ethambutol DST (18). Together with the numerous individual reports of low-level resistance and discordant phenotypic DST, these data illustrate low confidence in the capability of *emb*B Gly406Asp and Gly406Ala mutations to predict resistance at the current critical concentration.

To simulate improving concordance between genotypic and phenotypic DST in the current study, we applied lowered concentration thresholds to the study strains as shown in Table 3. This improved both the sensitivity and negative predictive values for ethambutol DST while maintaining high specificity and positive predictive value. A concentration of 2 μg/mL achieved the highest concordance.

**Table 3.** Predictive values for ethambutol critical concentrations applied to study strains.

| Critical Concentration (µg/mL) | PPV<sup>a</sup> (%) | NPV<sup>b</sup> (%) | Sensitivity (%) | Specificity (%) |
| --- | --- | --- | --- | --- |
| 5 | 100 | 45.0 | 25.0 | 100 |
| 4 | 100 | 50.0 | 37.5 | 100 |
| 3 | 100 | 66.7 | 68.8 | 100 |
| 2 | 100 | 100 | 100 | 100 |

<sup>a</sup>PPV, Positive Predictive Value. <sup>b</sup>NPV, Negative Predictive Value.

Before proposing a change to the critical concentration of ethambutol used for DST, we must consider the clinical relevance of low-level resistance. Denti *et al*. investigated the pharmacokinetics of ethambutol have reported *in vivo* plasma concentrations ranging from 2.7 – 4.8 μg/mL (46). Others have similarly reported serum concentrations of ethambutol under 5 μg/mL (47, 48). At the site of primary lesions in the lungs, however, ethambutol concentrations may be higher. Zimmerman *et al*. modeled ethambutol pharmacokinetics lungs of rabbits and found concentrations to be above 5 μg/mL, but these concentrations declined to below the critical concentration in the hours following treatment administration (49). As such, ethambutol concentrations below 5 μg/mL may be a biological reality for which low-level resistance is clinically relevant. Even so, we require a greater understanding of the clinical impact of low-level ethambutol resistance on ethambutol treatment outcomes.

While this study included all strains with *emb*B406 mutations in the Canadian National Reference Centre for Mycobacteriology culture collection 2003-2022, the limited number of strains included in this study must be acknowledged. Even so, our results corroborate a wealth of studies calling for improved concordance between molecular and phenotypic DST for ethambutol.

## Conclusion

We conclude that *emb*B406 mutations are associated with low-level ethambutol resistance. We predict that mutations outside of *emb*B, including *embR*, may promote variability in ethambutol resistance. Overall, this study supports the call to amend genotypic and phenotypic drug susceptibility testing to improve the sensitivity and concordance of ethambutol resistance predictions. Molecular DST is an invaluable tool to rapidly inform patient treatment regimens while gold culture methods remain time-consuming with long turnaround times. However, discordance between molecular and phenotypic methods must be acknowledged with caution. A greater understanding of the impact of *emb*B406 mutations on ethambutol resistance as well as the clinical significance of low-level resistance is required to inform improvement of DST.

## Data Availability

All raw sequence data generated in this work will be available upon publication within the NCBI Sequence Read Archive under BioProject ID PRJNA928676.

## Abbreviations

WHO: World Health Organization
TB: tuberculosis
MDR: multi-drug resistant
DST: drug susceptibility testing
WGS: whole genome sequencing
SNP: single nucleotide polymorphism
CLSI: Clinical Laboratory Standards Institute
FHA: forkhead-associated
LJ: Löwenstein-Jensen
MIC: minimum inhibitory concentration
PPV: positive predictive value
NPV: negative predictive value

